# A 50-gene high-risk profile predictive of COVID-19 and Idiopathic Pulmonary Fibrosis mortality originates from a genomic imbalance in monocyte and T-cell subsets that reverses in survivors with post-COVID-19 Interstitial Lung Disease

**DOI:** 10.1101/2023.10.22.563156

**Authors:** Bochra Tourki, Minxue Jia, Theodoros Karampitsakos, Iset M Vera, Alyssa Arsenault, Krystin Marlin, Carole Y Perrot, Dylan Allen, Forouzandeh Farsaei, David Rutenberg, Debabrata Bandyopadhyay, Ricardo Restrepo, Muhammad R. Qureshi, Kapilkumar Patel, Argyrios Tzouvelekis, Maria Kapetanaki, Brenda Juan-Guardela, Kami Kim, Panayiotis V Benos, Jose D. Herazo-Maya

**Affiliations:** Ubben Center for Pulmonary Fibrosis Research, Division of Pulmonary, Critical Care and Sleep Medicine. Department of Internal Medicine, Morsani College of Medicine, University of South Florida, Tampa, FL, USA; Department of Computational and Systems Biology, School of Medicine, University of Pittsburgh, Pittsburgh, PA, USA; Joint Carnegie Mellon - University of Pittsburgh Computational Biology Ph.D. Program, Pittsburgh, Pennsylvania, USA. Program in Computational Biology, Pittsburgh, PA, USA; Division of infectious Disease, Department of Internal Medicine, Morsani College of Medicine, University of South Florida, Tampa, FL, USA; Center for Advanced Lung Disease and Lung Transplant Program. Tampa General Hospital, FL, USA; Department of Respiratory Medicine, University of Patras, Patras, Greece; Department of Epidemiology, College of Public Health, and Health Professions, and College of Medicine, University of Florida, Gainesville, FL, USA

**Keywords:** 50 genes signature, COVID-19, high-risk, low-risk, post-Covid-19-ILD, IPF, resolution, 7Gene-M-MDSCs, CD4 T cells, CD8 T cells

## Abstract

**Background:** We aim to study the source of circulating immune cells expressing a 50-gene signature predictive of COVID-19 and IPF mortality.

**Methods:** Whole blood and Peripheral Blood Mononuclear cells (PBMC) were obtained from 231 subjects with COVID-19, post-COVID-19-ILD, IPF and controls. We measured the 50-gene signature (nCounter, Nanostring), interleukin 6 (IL6), interferon γ-induced protein (IP10), secreted phosphoprotein 1 (SPP1) and transforming growth factor beta (TGF-β) by Luminex. PCR was used to validate COVID-19 endotypes. For single-cell RNA sequencing (scRNA-seq) we used Chromium Controller (10X Genomics). For analysis we used the Scoring Algorithm of Molecular Subphenotypes (SAMS), Cell Ranger, Seurat, Propeller, Kaplan-Meier curves, CoxPH models, Two-way ANOVA, T-test, and Fisher’s exact.

**Results:** We identified three genomic risk profiles based on the 50-gene signature, and a subset of seven genes, associated with low, intermediate, or high-risk of mortality in COVID-19 with significant differences in IL6, IP10, SPP1 and TGFβ-1. scRNA-seq identified Monocytic-Myeloid-Derived Suppressive cells (M-MDSCs) expressing CD14^+^HLA DR^low^CD163^+^ and high levels of the 7-gene signature (7Gene-M-MDSC) in COVID-19. These cells were not observed in post-COVID-19-ILD or IPF. The 43-gene signature was mostly expressed in CD4 T and CD8 T cell subsets. Increased expression of the 43 gene signature was seen in T cell subsets from survivors with post-COVID-19-ILD. The expression of these genes remained low in IPF.

**Conclusion:** A 50-gene, high-risk profile in COVID-19 is characterized by a genomic imbalance in monocyte and T-cell subsets that reverses in survivors with post-COVID-19 Interstitial Lung Disease

## Introduction

The emergence and spread of 2019 coronavirus disease (COVID-19) led to an unparalleled, global public health crisis [1]. Despite major advances in the prevention and treatment of COVID-19, infections from emergent, severe respiratory syndrome coronavirus 2 (SARS-CoV-2) variants and long COVID still impose substantial burden on healthcare systems [2]. One of the most frequent manifestations of long COVID is post-COVID-19-Interstitial Lung Disease (Post-COVID-19-ILD), which is observed in a proportion of survivors from COVID-19 induced Acute Respiratory Distress Syndrome (ARDS) [3]. Recent evidence has demonstrated similarities between COVID-19-induced ARDS, post-COVID-19-ILD and IPF [4, 5], for example, both diseases are triggered by alveolar epithelial cell (AEC) injury, in COVID-19 the injury is viral and in IPF is unknown. AEC injury leads to recruitment of monocyte-derived alveolar macrophages, deregulated angiogenesis and vasculopathy, aberrant tissue remodeling and extracellular matrix deposition [5–7].

At the genomic level we have previously identified [8] and validated [9] a 50-gene signature predictive of IPF and COVID-19 survival [10] in circulating immune cells. In our previous work, we missed on the identification of additional COVID-19 endotypes based on the 50-gene signature, we lacked correlation with cytokine data, and we did not evaluate the clinical applicability of our genomic risk profiles by a test that could be widely used in clinical practice. Finally, we did not perform in-depth immunophenotyping in patients with COVID-19, post-COVID-19-ILD and IPF. In the present work, we analyzed over 216 hospitalized patients with COVID-19 and identified the presence of three genomic risk profiles based on the 50-gene signature (and a subset of these genes) associated with a low, intermediate, or high-risk of mortality with significant differences in pro-inflammatory and pro-fibrotic cytokines.

We also used single-cell RNA sequencing to study genes with increased (seven genes) and decreased expression (43 genes) in circulating immune cells associated with increased risk of mortality in COVID-19 and discovered a novel subtype of Monocytic-Myeloid-Derived Suppressive (M-MDSC) cells expressing CD14^+^HLA-DR^low^CD163^+^MCEMP1^+^PLBD1^+^S100A12^+^TPST1^+^IL1R2^+^FLT3^+^HP^+^ responsible for the high-risk genomic profile. We denominated these cells as 7Gene-M-MDSC. These cells were not observed in survivors with post-COVID-19-ILD and in IPF patients in our cohort. Our findings suggest that a 50-gene, high-risk profile may represent the imbalance between increased 7GeneM-MDSC and decrease CD4 T and CD8 T subsets expressing the 43-gene signature in patients with increased risk of COVID-19 mortality. While increased expression of the 43-gene signature was seen in T cell subsets from survivors with post-COVID-19-ILD, the expression of these genes remained low in IPF.

## Methods

### Human specimens and study design

Whole blood and Peripheral Blood Mononuclear cells (PBMC) were obtained from 216 hospitalized patients with COVID-19 from the University of South Florida (USF)/Tampa General Hospital (TGH). Cohorts were split in three based on time of collection and experiments performed. For the 50-gene signature analysis, whole blood samples were collected from 75 patients recruited between 07/2021 and 02/2022 at USF/TGH within two days (±2) from hospital admission. For time course analyses, 23 and 14 blood samples were collected at days six (±2.3) and 13 (±1.5), respectively, from hospital admission (Cohort one, **Fig1A**). For the 7-gene signature analysis, PBMC were collected from 141 patients admitted to USF/TGH between 06/2020 to 10/2020 within 3.7 days from hospital admission (Cohort two, **Fig1I**). For single-cell RNA sequencing, we included three hospitalized patients with COVID-19, five patients with post-COVID-19-ILD, six patients with IPF and four controls (Cohort three, **Fig2A**). Post-COVID-19-ILD patients were enrolled with an average of 7 months (±5) from COVID-19 diagnosis. These patients developed pulmonary sequalae defined as clinical symptoms, abnormal radiographic findings of Interstitial Lung Disease and Diffusion of the Lung for Carbon Monoxide (DLCO) less than 70. All studies were approved by Institutional Review Boards (Pro00032158 and Study00085). Clinical data were recorded for each patient at the time of admission and during time course studies.

**Figure 1.**
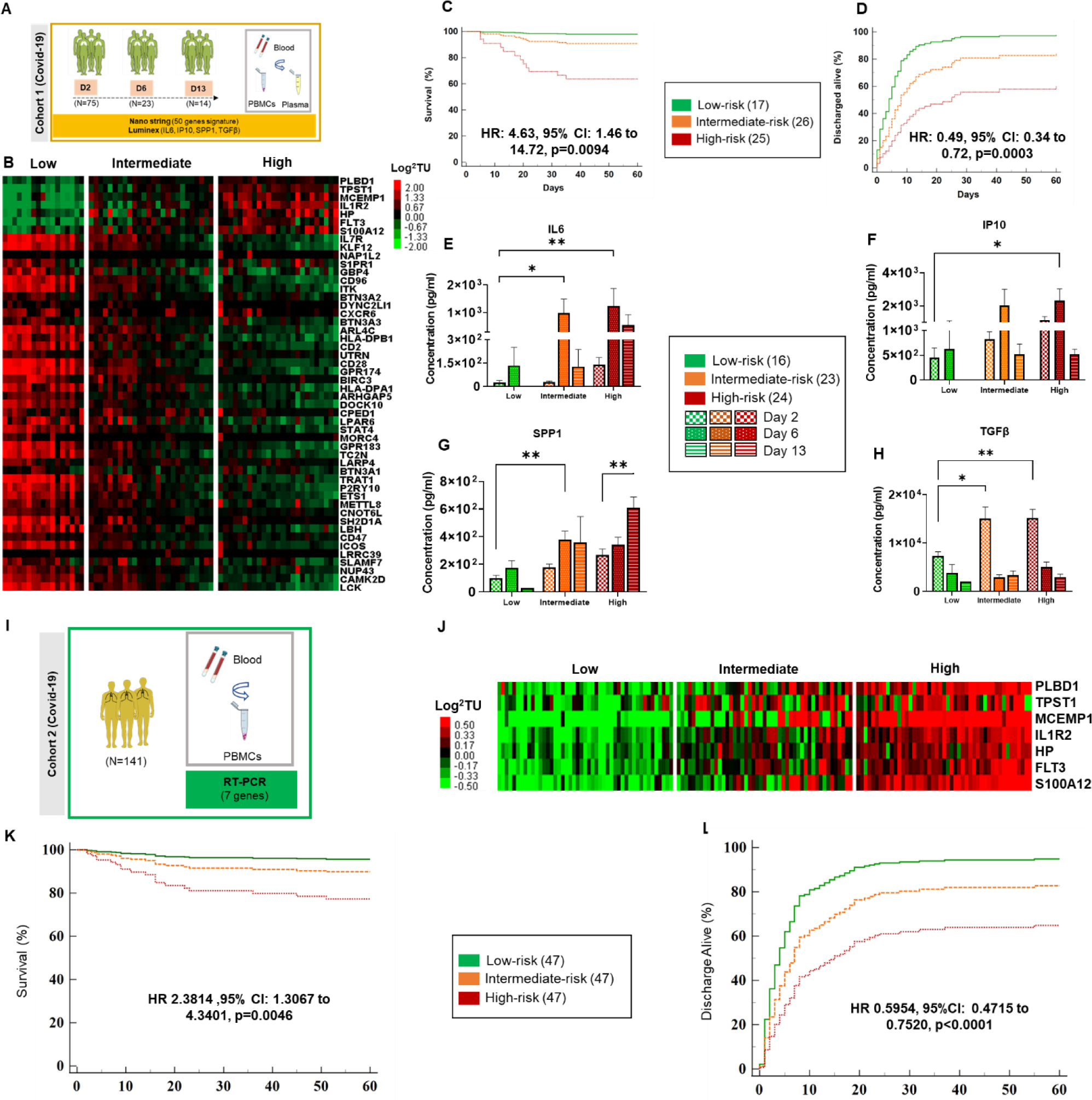
A 50-gene signature can be used to identify three molecular endotypes associated with differences in COVID-19 survival and cytokine profiles. **A**. Study design of the 50-gene signature and cytokine analysis in COVID-19 patients (Cohort 1). **B.** Heatmap of COVID-19 patients based on the 50-gene signature discriminates three risk groups (low, intermediate, and high) based on SAMS. Every column represents a patient, and every row represents a gene. Log-based two-color scale is adjacent to the heatmap. Red denotes increased expression and green denotes decreased expression. Gene expression data is represented as Log2 normalized expression values. **C-D.** Time to death and time to discharge by day 60 in hospitalized patients with COVID-19, respectively. **E-H.** Plasma cytokine concentrations (IL6, IP-10, SPP1 and TGFβ) in low, intermediate, and high-risk profile patients with COVID-19 at days 2, 6 and 13 post admission. The data is presented as an average of triplicate values ± SEM for each group. Two-way ANOVA test (GraphPad software) Tukey’s multiple comparisons were used; * p<0.05. **I.** Study design of 7-gene signature analysis by RT-qPCR in PBMCs from COVID-19 patients (Cohort 2). **J.** Heatmap of COVID-19 patients based on the 50-gene signature discriminates three risk groups (low, intermediate, and high) based on SAMS Up scores. Heatmap nomenclature is the same as in Figure 1A. **K-L.** Time to death and time to discharge by day 60 in hospitalized patients with COVID-19 respectively in cohort two. The data is presented as an average of triplicated TUs values ± SEM for each group. * p<0.05

### nCounter analysis system (Nanostring) experiments

A custom code set including the 50-gene signature was generated using the nCounter analysis system as previously described [9]. Briefly, 200 ng of total RNA was extracted using Pax gene blood miRNA kit (Cat#763134, PreAnalytix, Qiagen). RNA quality control was confirmed by nanodrop and tape station 4150 (Agilent). Samples were run in batches of 12 samples. RCC raw files were generated by nCounter system and analyzed by nSolver 4.0 software. Data were normalized to the geomean of seven housekeeping genes (*GUSB, GAPDH, TRAP1, FPGS, ACTB, DECR1, FARP1*). Data was log transformed and presented as Log2 and used to calculate the Scoring Algorithm for Molecular Subphenotypes (SAMS) [9, 10] to stratify risk profiles as previously described.

### Luminex

We measured cytokine concentrations of 121 plasma samples from COVID-19 patients from Cohort 1 using a customized, Bioplex 200 compatible, human cytokine panel including, IL6, IP10, SPP1 and TGFβ-1 (#FCSTM18-06, R&D Systems). The equipment was calibrated (Cat# 171203060, BIO-RAD) and validated (Cat#171203001, BIO-RAD) prior utilization.

### Taqman RT-qPCR

141 blood samples were collected from hospitalized COVID-19 patients at USF/TGH from Cohort 2. To perform RT-qPCR experiments (Quantstudio6), we used qPCR kit, SuperScript IV VILO Mater Mix (Cat#11756050), 10 TaqMan probes, MCEMP1 (Cat #Hs00545333_g1), PLBD1 (Cat #Hs00227344_m1), IL1R2 (Cat #Hs00174759_m1), HP (Cat #Hs00978377_m1), FLT3 (Cat#Hs00174690_m1), TPST1 (Cat#Hs01041471_m1), S100A12 (Cat #Hs00942835_g1), ACTB (Cat#Hs99999903_m1), B2M (Cat# Hs00187842_m1) and RPS18 (Cat# Hs01375212_g1) (Thermofisher). Triplicated CTs values were geo-normalized to reference genes. Data was represented as transcript unit (TU, 2^^−10 (ΔCT)^). SAMS was used to identify genomic risk profiles based on the 7-gene signature.

### Single-cell RNA sequencing (scRNA-seq) experiments

For scRNA-seq, we used cryopreserved PBMC from healthy controls and patients with COVID-19 (from Cohort 1), post-COVID-19-ILD and IPF as described above. IPF diagnosis was based on current guidelines [11]. A single-cell suspension from PBMC from each patient was quantified and analyzed for viability using the Cell counter 3 (Countess 3, Invitrogen) and then loaded onto the 10X Genomics Chromium Single Cell Controller for isolation of single cells (10X Genomics). Briefly, 5000-6000 PBMC from each of the samples were targeted for recovery. The single cells, reagents, and 10x Genomics gel beads were encapsulated into individual nanoliter-sized Gel beads in Emulsion (GEMs) and then reverse transcription (RT) of poly-adenylated mRNA was performed inside each droplet. Post GEM-RT Cleanup and cDNA was amplified, purified, and cDNA libraries were then prepared in bulk reactions using the Chromium Next GEM Single Cell 3ʹ Kit v3.1 Library Prep Kit. From sequencing, approximately 35000 mean reads per cell were generated on the Illumina NextSeq. FASTQ files were generated further demultiplexing, barcode processing, alignment, and gene counting steps for analysis.

### scRNA-seq analysis and quality control

scRNA-seq feature count matrices were constructed using Cell Ranger (v7.1.0), aligning reads to the GRCh38 2020 reference genome. Subsequent quality control and data processing were performed with the Seurat package (v4.3.0) [12]. Cells with less than 200 detected genes were discarded, as well as cells with more than 15% mitochondrial genes. DropletUtils (v1.14.2) identified empty droplets [13], while doublets were detected using Scrublet (v0.2.3-0) [14]. DecontX was used to detect ambient RNA contamination [15]. We excluded identified empty droplets, doublets, and those with a contamination score above 0.2, alongside cells with more than 9 UMIs mapping to the hemoglobin subunit beta (HBB) gene (representing red blood cells) for further analysis. For cell type annotation, we used SCTransform for data normalization and identification of variable features. This was followed by principal component analysis (PCA) and batch effect correction using Harmony [16]. Graph-based clustering and UMAP embedding were generated based on Harmony embeddings. Cell types were assigned to each cluster guided by canonical marker gene expression [17–19]. For cell type proportions analysis, we normalized the number of cells within a given cell type by the total number of cells per subject. For Differentially Expressed Gene (DEG) analysis, we used the Seurat software [20] (v4.3.0) “FindMarkers” function to identify differentially expressed DEGs across disease groups. Differential abundance analysis was performed for each cell type across different disease conditions, utilizing the ‘propeller’ method [21] as integrated within the ‘speckle’ R package.

### Statistics

The Kaplan–Meier method for survival and time of discharge analysis was used for the nCounter data and RT-qPCR data (Cohorts 1 and 2) using MedCalc version (v12.104). Cumulative incidence curves were analyzed among the three groups. Cox proportional-hazards models were used to estimate the hazard ratio and 95% confidence interval. Two-way ANOVA followed by Tukey multiple comparisons test by Graphpad prism software was used for the analysis of cytokines levels. T-test, Fisher’s exact test was used to compare cell proportions and clinical data respectively across different disease groups. The Wilcoxon rank test was used to compare DEG analysis within each group.

## Results

### A 50-Gene signature can be used to identify three molecular endotypes associated with differences in COVID-19 survival and cytokine profiles

SAMS identified three risk-profile groups of COVID-19 patients (low, intermediate, and high-risk) based on the 50-gene signature in Cohort 1. A 0.91 up score and −3.42 down score were used to identify the high-risk group. Intermediate and low-risk groups were then split based on a 0.15 up score and a −0.15 down score (**Fig1A-B**). These three risk-profile groups had significant differences in mortality (HR: 4.63, 95% CI: 1.46 to 14.72, p=0.0094) and time to discharge (HR: 0.49, 95% CI: 0.34 to 0.72, p=0.0003) after adjustment for Charlson Comorbidity Index (**Fig1 C-D**). Table 1 summarizes the clinical characteristics of patients in this cohort.

**Table 1.** Demographics and clinical data of cohort 1.

| Risk profile | N | Age | Gender Predominant | CRP | Ferritin |
| --- | --- | --- | --- | --- | --- |
| High-Risk | 25 | 55.5±15.7 | 14 Males | 12.0±8.6 | 1816.9±1340.9 |
| Intermediate-Risk | 26 | 51.6±15.8 | 15 Males | 11.3±8.7 | 1902.8±2022.9 |
| Low-Risk | 17 | 57.3±18.4 | 9 Females | 3.4±4.1 | 491.4±521.1 |
| P value | NA | NS | NA | 0.006 | 0.03 |

We then investigated whether 50-gene risk profiles were associated with temporal changes in proinflammatory and profibrotic cytokines including IL6, IP10, SPP1 and TGF-β at baseline and over time in COVID-19. While we did not find significant difference in IL6, IP10, SPP1 levels between the three risk groups at baseline (**Fig1E-H**), TGF-β levels were significantly increased in the high-risk group compared to the low-risk group (15,193.17 ± 8,214.71 vs 7,315.926 ± 3,467.94pg/ml p<0.01). Our results also demonstrated an increase of IL6, IP10 and SPP1 at day6 compared to baseline independent of the 50-gene risk profile. 50-gene, high-risk profile patients displayed the highest levels of IL6 at day6 compared to intermediate and low-risk patients, respectively (1,236 ± 8.87 vs 980 ± 34.69 vs 134± 116 pg/ml, p<0.01) (**Fig1E**). The same trend of cytokine levels was noted with IP10 at day6 compared to intermediate and low-risk group patients, respectively (2,323 ± 699 vs 2,029 ± 980 vs 625± 470 pg/ml, p<0.01) (**Fig1F**) and similar results were noted with SPP1 (**Fig1G**). Most of the studied cytokines trended down over time except for SPP1 which increased at day 13 in patients with a 50-gene, high-risk profile (**Fig1-G).** In terms of TGF-β, we found the highest levels in day2 in the high-risk (1,5193 ± 1,712 pg/ml) and intermediate-risk (1,5035 ± 2222 pg/ml) groups compared to the low-risk group (7315± 723 pg/ml, p<0.01) (**Fig1H**). We observed a drastic decrease of circulating TGFβ on day6 in the high-risk (threefold change) and intermediate-risk (fivefold change) groups respectively compared to baseline. No significant changes in the levels of studied cytokines were observed in the low-risk group over time (**Fig1H**).

Finally, we measured the expressions of *MCEMP1*, *PLBD1*, *TPST1*, *S100A12*, *IL1R2*, *HP* and *FLT3* by RT-qPCR in Cohort 2 patients (**Fig1I**) and calculated SAMS UP scores. The three risk groups were split by the Up score based on tertiles and depicted on a heatmap (**Fig1J**). Our 7-gene RT-qPCR test confirmed the presence of three endotypes of COVID-19 with significant differences in mortality (HR 2.3814, 95% CI: 1.3067 to 4.3401, p=0.0046) (**Fig1K**) and time to discharge alive (0.5954, 95%CI: 0.4715 to 0.7520, p<0.0001) (**Fig1L**) after adjustment for Charlson Comorbidity Index. Table 2 summarizes the clinical characteristics of patients in this cohort. In summary, we demonstrated the presence of three genomic risk profiles of COVID-19 patients with significant differences in survival and cytokine profiles by two different methods and in two separate cohorts.

**Table 2.** Demographics and clinical data of cohort 2.

| Risk profile | N | Age | Gender Predominant | CRP | Ferritin |
| --- | --- | --- | --- | --- | --- |
| High-Risk | 47 | 60.06±19.09 | 29 Males | 15.04±65.60 | 2463.6±534.2 |
| Intermediate-Risk | 47 | 59.23±12.51 | 27 Males | 13.21±20.50 | 1788.8±3449.3 |
| Low-Risk | 47 | 59.28±18.80 | 25 Females | 11.12±8.93 | 477.4±534.3 |
| P value | NA | NS | NA | NS | NS |

### The 7-gene signature predictive of COVID-19 mortality can be identified in a novel subtype of Monocytic-Myeloid Derived Suppressive Cells

To delineate the cellular source of the 7-gene signature that predicts COVID-19 mortality when overexpressed (MCEMP1, PLBD1, S100A12, FLT3, TPST1, IL1R2 and HP), we performed scRNA-seq from frozen PBMC of healthy controls, COVID-19, post-COVID-19-ILD and IPF patients (**Fig2A, Table 3**). All immune cell clusters (**Fig2B**) were identified based on the expression levels of different markers by reference to COVID-19 [22] and IPF atlas [23, 24]. In terms of cell frequencies, we noticed a significant increase in CD14^+^CD163^+^HLA-DR^low^ monocytes, platelets and plasmablasts, and a decrease in naive CD4T, memory CD8T GZMB^+^ and dendritic cells when comparing COVID-19 versus post-COVID-19-ILD. Notably, we identified a significant increase in Hematopoietic and Progenitor Stem Cells (HSPC), dendritic cells and plasmablasts when comparing post-COVID-19-ILD with IPF patients (**Fig2C**) (**Table 4, Supplementary file 1**). Among the four conditions studied, the expression of the seven gene signature was limited to circulating monocytes and platelets (to a lesser degree) compared to other immune cell populations (**Fig2D).** All seven genes were highly expressed in all the COVID-19 patients and in one IPF patient (**Fig2E**). Out of the seven genes, three genes (*MCEMP1, PLBD1* and *S100A12*) were expressed in monocytes in the four groups studied. In post-COVID-19-ILD samples, all seven genes had decreased expression compared to COVID-19. (**Fig2E, Supplementary file 2**). To gain more insight on the expression of the seven genes of interest in monocyte subtypes, we employed uniform manifold approximation and projection (UMAP) of these cells to connect them with integrated immune signatures in the subgroups studied (**Fig2F**). Monocyte subpopulations were characterized by the expression of CD45, CD14, CD16, HL-DRA, CD163, CD11b, CD11c, S100A12 and S100A8 (**Fig2G**). Out of the three classical monocyte subpopulations identified: HLA-DR^hi^CD163^−^, HLA-DR^low^CD163^−^ and HLA-DR^low^CD163^+^ (**Fig2F),** two were classical monocytes HLA-DR^low^ expressing CD33^+^ and CD15^−^ (FUT4) qualifying as Monocytic-Myeloid Derived Suppressive Cells (M-MDSCs), one of them (HLA-DR^low^CD163^+^), expressed exclusively in COVID-19 patients (**Fig2F, Supplementary figure 2).**

**Figure 2.**
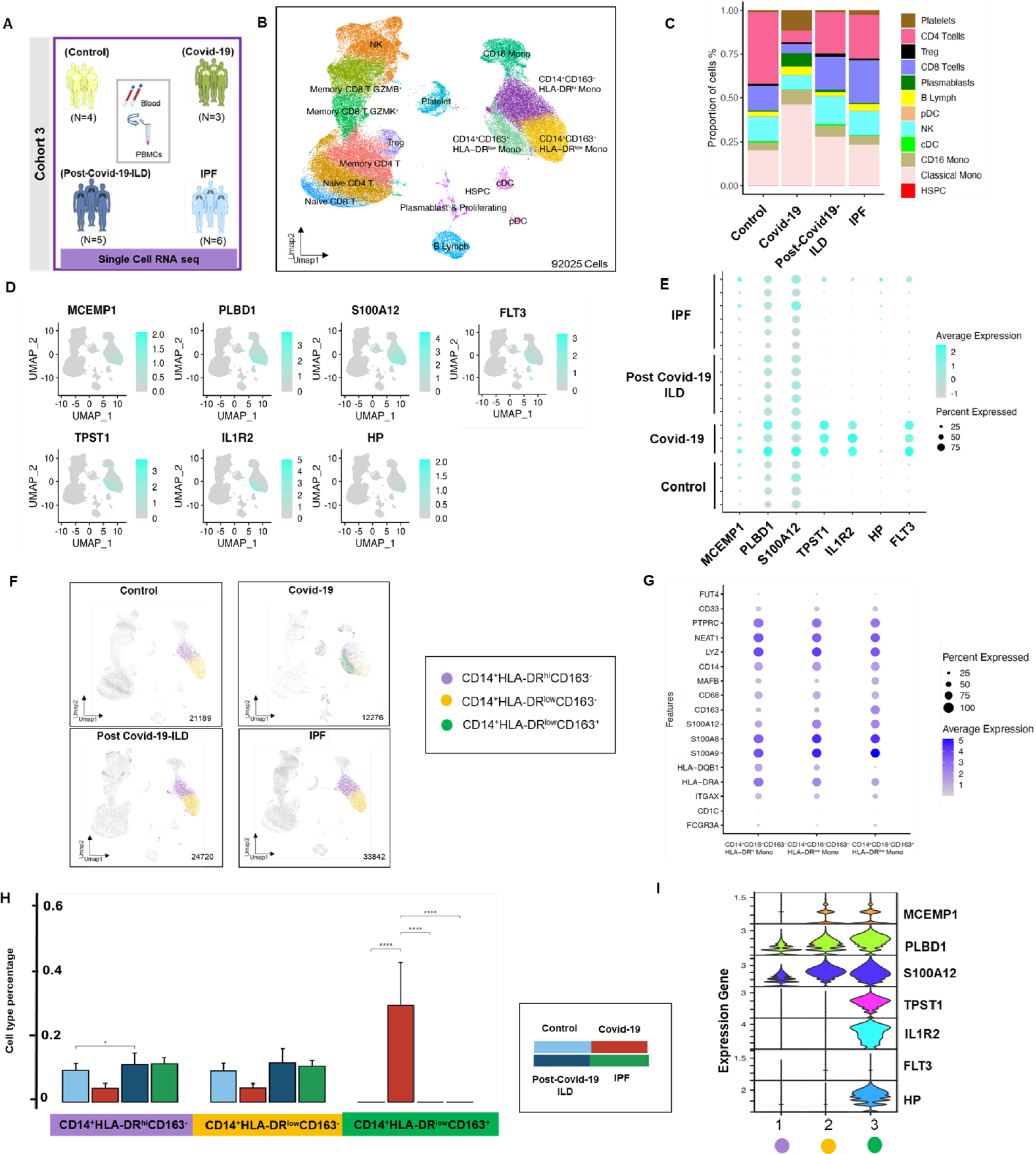
Increased expression of seven genes associated with increased risk of mortality in COVID-19 can be identified in a novel subtype of Monocytic-Myeloid Derived Suppressive Cells. **A.** Study design of scRNA-seq in cohort three**. B.** Uniform manifold approximation and projection (UMAP) embedding plots of 92027 single-cells of the four studied conditions: controls, COVID-19, post-COVID-19-ILD and IPF patients, showing the cellular landscape with cluster-colored annotations. **C.** Stacked bar graph of cell count percentage of immune cells in each condition. **D**. Aggregated UMAPs of the four studied conditions projecting the major expression of each gene of the 7-gene signature: MCEMP1, PLBD1, S100A12, FLT3, TPST1, IL1R2, HP on immune cells (aqua blue color). **E.** Dot plot of seven increased genes in high-risk patients across controls, COVID-19, post-COVID-19-ILD and IPF. **F.** UMAPs of 21189 cells from four controls patients, 12276 cells from three COVID-19, 24720 cells from five post-COVID-19 and 33842 from six IPF patients were analyzed and integrated in four separate UMAPs to represent three monocyte subpopulations, grouped in a color-coded manner. **G.** Dot plots comparing expression of 15 selected marker genes for clustering classical monocytes populations (CD14^+^CD16^−^). Three subpopulations of classical monocytes were identified based on the expression of HLA-DR and CD163 markers. Dot size is proportional to the percentage of cells expressing the gene in each subcluster. Color intensity is proportional to the average scaled log2-normalized expression within the subcluster. **H.** Bar graphs of cell percentages in the three classical monocyte subpopulations identified (CD14^+^CD16^−^) HLA-DR^hi^CD163^−^, HLA-DR^low^CD163^−^ and HLA-DR^low^CD163^+^, stratified by conditions. **I.** Violin plot of the seven gene expressions on the three classical monocyte subgroups identified in panel **H**. Data are presented scale/log normalized as average expression of all cells within a given group. The propeller method and T test were used to compare cell frequencies in each group. * p < 0.05, ** p < 0.01, *** p< 0.001, and **** p < 0.0001.

**Table 3.**
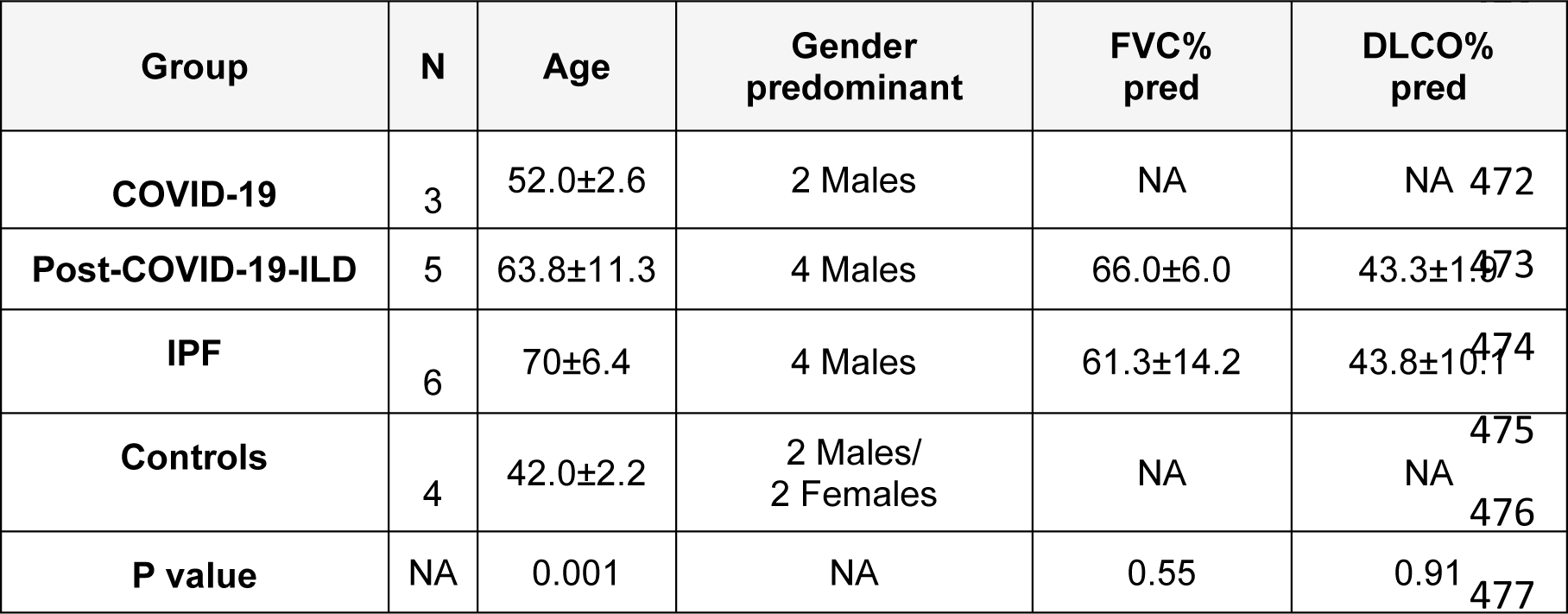
Demographics and clinical data of cohort 3.

**Table 4.**
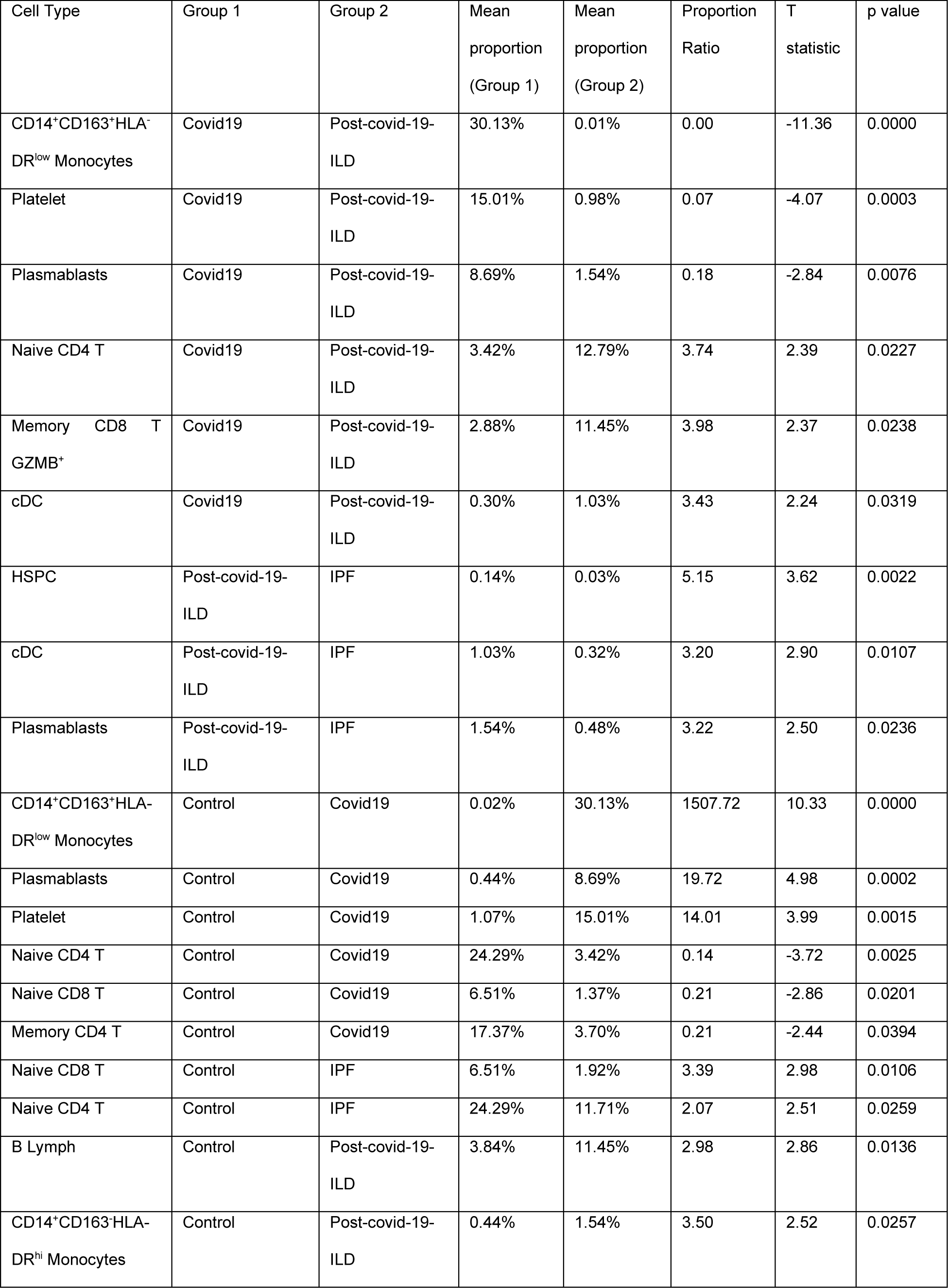
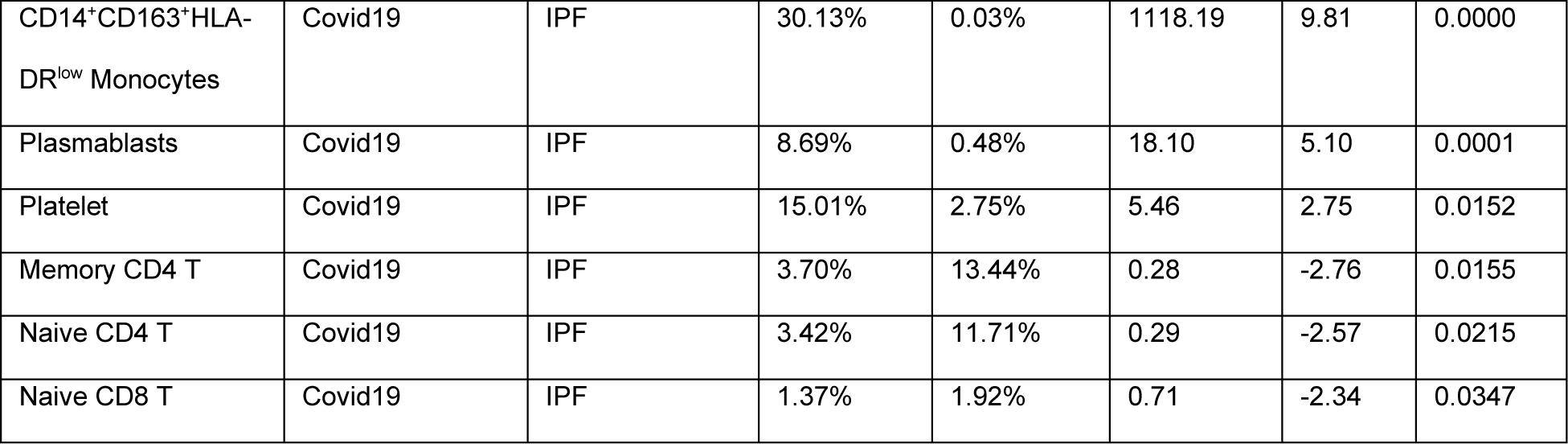
Cell frequencies among studied groups and abundance analysis. Only cell frequencies with P<0.05 are shown.

Cellular composition of the three monocyte subpopulations varied among controls, COVID-19, post-COVID-19-ILD and IPF. CD14^+^HLA-DR^low^CD163^+^ M-MDSCs were exclusively expressed in COVID-19 (0.36) and absent in the three other groups (**Fig2H).** The COVID-19 group displayed low percentages of HLA-DR^hi^CD163^−^ (0.05) and HLA-DR^low^CD163^−^ (0.05) cells compared to control group (0.10 and 0.10, respectively). No significant difference was observed between these HLA-DR^hi^CD163^−^ and HLA-DR^low^CD163^−^ monocytes in post-COVID-19-ILD (0.13 and 0.14) and IPF (0.12 and 0.11) patients compared to controls. An increase in HLA-DR^hi^CD163^−^ monocytes was observed in post-COVID-19-ILD patients compared to controls (0.13 vs 0.10) (**Fig2H**). Finally, we performed deep immune profiling and studied the expression of the seven gene signature in the three classical monocytes subpopulations identified. Increased expression of the seven genes was observed in CD14^+^HLA-DR^low^CD163^+^ cells compared to CD14^+^HLA-DR^hi^CD163^−^ and CD14^+^HLA-DR^low^CD163^−^ cells (**Fig2I**). Taken together, our results suggests that CD14^+^HLA-DR^low^CD163^+^ M-MDSCs are the cellular source of a high-risk genomic profile characterized by increased expression of *MCEMP1, PLBD1, S100A12, FLT3, TPST1, IL1R2* and *HP* and associated with increased risk of mortality in COVID-19 thus we denominated these cells as 7Gene-M-MDSCs.

### The 43-gene signature originates from CD4 T, and CD8 T cell subsets and its expression increases in post-COVID-19-ILD

To determine the cellular source of the 43 genes with decreased expression in high-risk COVID-19 patients, we analyzed our scRNA-seq dataset with integrated DEG analysis. To functionally map the expression of the 43-genes on T cells across disease groups, we plot six clusters of T cells in a color-coded manner, stratified by conditions. Most of the genes of the 43 gene signature are expressed in Tregs, memory CD4 T, memory CD8 T GZMK^+^, naive CD4T, naive CD8 T, memory CD8 T GZMB^+^ (**Fig3A**). Overall, we identified a low transcriptomic signal in lymphocytes in COVID-19 compared to the other conditions studied (**Fig3B**) which is reflected by a significantly lower percentages of memory CD4 T, Naive CD4 T, Naive CD8 T and Memory CD8 T GZMB+ cells in COVID-19 compared to controls, IPF and post-COVID-19-ILD (**Fig3C**). Notably, we did not identify significant changes in the composition of the T cell compartment between post-COVID-19 and IPF.

**Figure 3.**
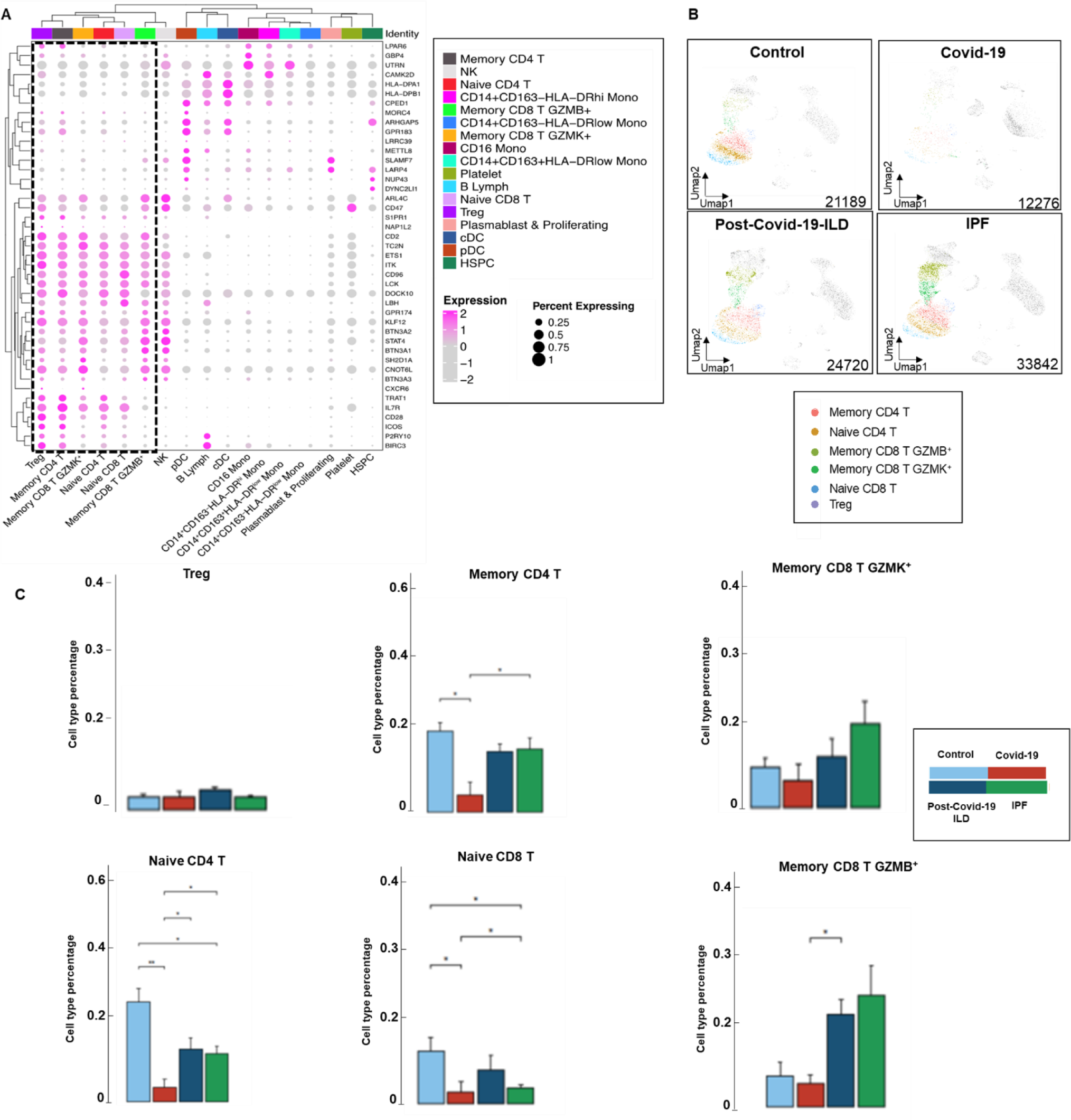
CD4 T and CD8 T cell subsets are the main source of the 43-gene signature. **A. Clustered** dot plot of the 43 genes signature in all aggregated groups (controls, COVID-19, post-COVID-19-ILD and IPF patients) in each identified cell cluster. **B.** Separate UMAPs representing T immune cells subpopulations distributions in controls, COVID-19, post-COVID-19-ILD and IPF patients in a color-coded manner. Data are presented scale/log normalized as average expression of all cells within a given group. **C.** Bar graphs of T-cell subset percentages stratified per conditions. The propeller method and T test were used to compare cell frequencies in each group. * p < 0.05, ** p < 0.01, *** p< 0.001, and **** p < 0.0001

When we looked at gene expression changes of genes of the 43-gene signature, we noticed that survivors with post-COVID-19-ILD had overall increased expression of the 43 genes of interest, compared to COVID-19 and IPF patients. When compared to COVID-19, post-COVID-19-ILD patients had increased expression of most of the genes of the 43-gene signature in naive CD4 T and memory CD4 T cells, Tregs, memory CD8 T GZMB^+^, memory CD8 T GZMK^+^ and naive CD8 T cells (**Fig4A-F**). To better represent the overall changes in gene expression of the 43-gene signature, we calculated the median of the average log2fold change values across the 43 genes, in each T cell subtype, in the four conditions studied (post COVID-19-ILD versus COVID-19 and, IPF versus post COVID-19-ILD) (**Table 5 and supplementary file 1**). Positive values represent overall higher expression and negative values represent overall lower expression of the 43-gene signature in the first group, respectively. When comparing post-COVID-19-ILD with COVID-19 patients, we found a median of the average log2fold change values that ranged between 0.1 and 0.57 indicating that post-COVID-19-LD patients had overall increased expression of the 43 gene signature in each T cell subtype. The most pronounced effect was seen with increased expression of these genes in memory CD8 T GZMK^+^ cells with a median of the average log2fold change values of 0.52. In this subgroup of CD8 T cells, we found 26 DEG out of 43 genes between COVID-19-ILD versus COVID-19 (Bonferroni adjusted P<0.05). When comparing IPF with post-COVID-19-ILD patients, we found a median of the average log2fold change values that ranged between −0.16 and −0.38 indicating that IPF patients had overall decreased expression of the 43 gene signature in each T cell subtype. The most pronounced effect was seen with decreased expression of these genes in naive and memory CD4 T cells (median of the average log2fold change of −0.38 and −0.35, respectively). In naive CD4 T cells, we found 28 DEG out of 43 genes (Bonferroni adjusted P<0.05) and in memory CD4 T cells we found 34 DEG out of 43 genes between IPF versus COVID-19 (Bonferroni adjusted P<0.05). In summary, our results demonstrate that decreased expression of the 43-gene signature in COVID-19 and in IPF originates from CD4 T and CD8 T cell subsets, a finding that reverses in post-COVID-19-ILD.

**Figure 4.**
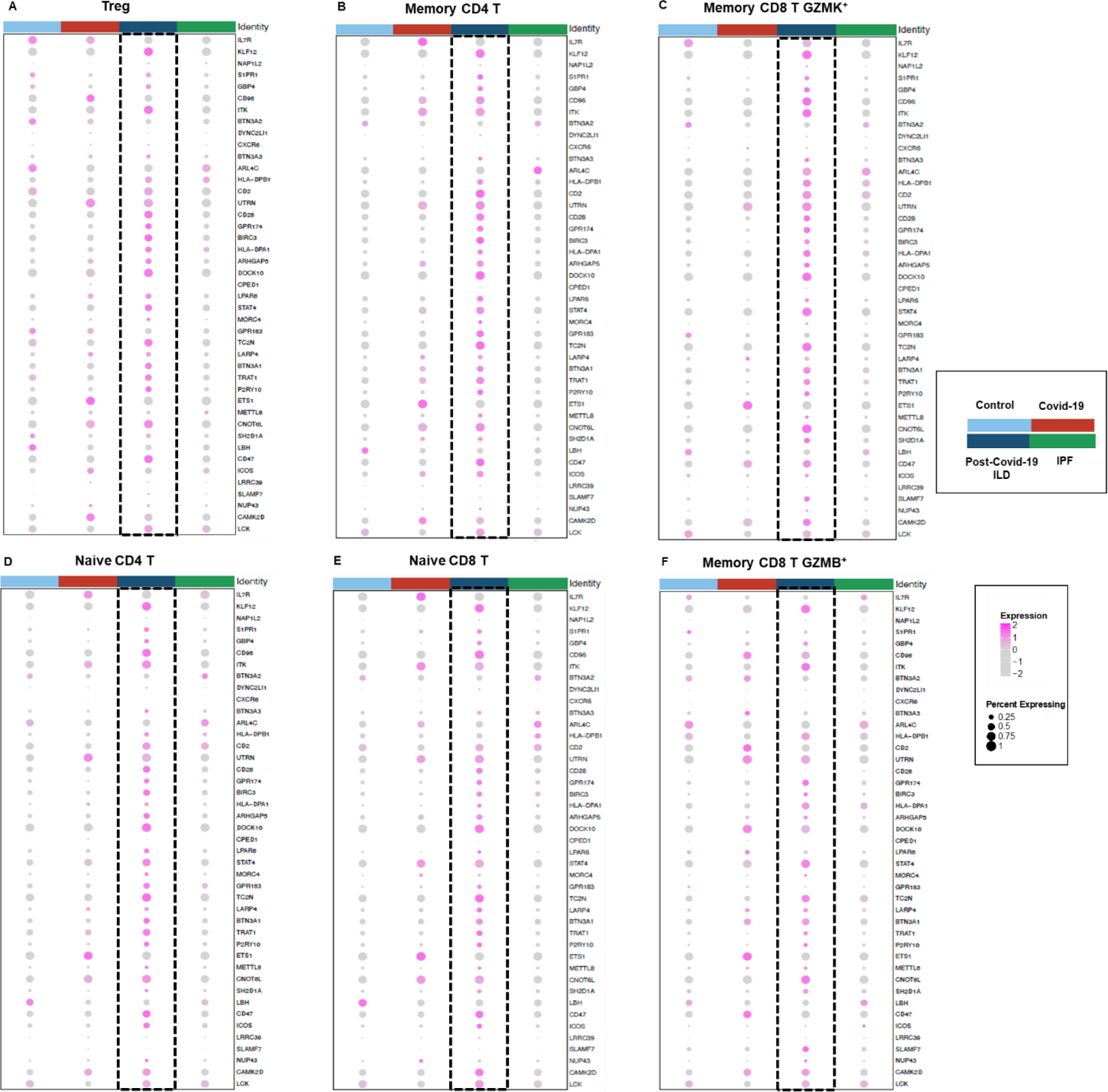
Resurgence of the 43-gene signature in survivors with post-COVID-19-ILD. **A.** Dot plot of genes of the 43 gene signature in Tregs, memory CD4 T cells, memory CD8 T GZMK^+^, naive CD4 T, naive CD8 T and memory CD8 T GZMB^+^ cells, respectively. Dot size is proportional to the percentage of cells expressing the gene in each subcluster. Color intensity is proportional to the average scaled, log-normalized expression within the disease group. Data are represented as average of log2. Log-based two-color scale is adjacent to the dot-plots.

**Table 5.** Summary statistics of DEG in scRNA-seq between post COVID-19-ILD versus COVID19 and IPF versus post COVID-19-ILD. *Average log2 fold change. Positive values indicate that the gene is more highly expressed in the cell cluster. The median of average log2 fold change was calculated using the average log2-fold change values of the 43 genes in each comparison. ** Bonferroni corrected P value

|  | Post COVID19-ILD Vs COVID19 |  |  | IPF Vs post COVID-19-ILD |  |  |
| --- | --- | --- | --- | --- | --- | --- |
| Cell types | Median of average log2fold change in 43 genes* | #43 Genes with P<0.05 | #43 Genes with p adjusted <0.05** | Median of average log2-fold change in 43 genes | #43 Genes with P<0.05 | #43 Genes with p adjusted <0.05 |
| Naive CD4 T | 0.27 | 35 | 20 | -0.38 | 37 | 28 |
| Memory CD4 T | 0.10 | 26 | 15 | -0.35 | 41 | 34 |
| Treg | 0.21 | 19 | 5 | -0.24 | 29 | 10 |
| Memory CD8 T GZMB+ | 0.13 | 25 | 15 | -0.16 | 37 | 26 |
| Memory CD8 T GZMK+ | 0.57 | 35 | 26 | -0.29 | 37 | 26 |
| Naïve CD8T | 0.18 | 26 | 7 | -0.19 | 28 | 13 |

## Discussion

In this study, we aimed to validate the performance of a 50-gene signature (and a subset of seven of these genes) previously shown to predict IPF [8, 9] and COVID-19 mortality [10]. We also aimed to identify the cellular source of these gene expression changes and to analyze the expression of these genes at the single-cell level through the course of normal health, acute COVID-19 infection, post COVID-19 infection with pulmonary fibrosis and pulmonary fibrosis without infection. We have previously identified a 50-gene signature that was able to discriminate two groups of patients with COVID-19 with increased risk of mortality and poor disease outcomes [10]. In the present study, we were able to discriminate three risk profiles (low, intermediate, and high risk) based on the 50-gene signature with significant differences in outcomes and cytokine profiles. To study the clinical applicability of a genomic test based on genes of the 50-gene signature, we designed a RT-qPCR panel including seven of these genes. Strikingly, we were able to validate the existence of three risk profiles of COVID-19 patients with increased risk of mortality and poor outcomes in a separate cohort when using the 7-gene RT-qPCR test. A 7-gene RT-qPCR test is a reliable, non-labor intense, fast and inexpensive way to predict mortality [25] [26]. Our study also focused on identifying the cellular source of these gene expression changes using scRNA-seq. We found that 7Gene-M-MDSCs were present almost exclusively in COVID-19 patients with severe disease. When we studied the expression of genes of the 43-gene signature at the single-cell level, we identified their expression mostly in naive CD4 T and memory CD4 T cells, Tregs, memory CD8 T GZMB^+^, memory CD8 T GZMK^+^ and naive CD8 T cells. Interestingly, the expression of these genes was higher in survivors with post-COVID-19-ILD, but they remained low in IPF which is a thought-provoking finding because most patients with post-COVID-19-ILD have partial or complete resolution of their pulmonary fibrosis while most patients with IPF have disease progression.

We have previously demonstrated the presence of a large subset of IPF patients with a high-risk genomic profile based on the 52-gene signature and increased monocyte counts [27] both associated with increased mortality [9] suggesting that 7Gene-M-MDSCs exists in IPF, but we may have missed their detection in this study due to the limited number of IPF samples analyzed. Also, it is possible that patients with post-COVID-ILD who do not recover, progress or have persistently long COVID symptoms, may also have a high number of 7Gene-M-MDSCs, driving a state of persistent immune dysregulation. Several lines of evidence have shown that COVID-19 is associated with dysregulated myeloid cell compartment [19, 28–31]. Myeloid cells have been found highly and aberrantly activated in COVID-19 [28, 32], with dysfunctional HLA-DR^lo^CD163^hi^ and HLA-DR^lo^S100A^hi^ CD14^+^ monocytes being present in patients with severe disease [33]. Lymphopenia has been an established negative prognostic marker in COVID-19 [34], while T cell subpopulations of patients with COVID-19 have exhaustion features [35–37]; yet, this is still a matter of debate [37, 38]. Importantly, single-cell bronchoalveolar lavage transcriptomic profiling comparing post-COVID-19-ILD patients with inflammatory and ‘’fibrotic-like’’ changes, showed more abundant expression of CD4 central memory and CD8 effector memory T cells in the inflammatory arm, suggesting a faster immune response recovery in patients with less pronounced radiographic abnormalities [39]. Our study further expands on the major role of T cell recovery in post-COVID-19-ILD and T-cell exhaustion in IPF [40] (**Figure 5**). Despite the reproducibility and relevance of our findings, we acknowledge the limitations of our study. First, we did not investigate the underlying mechanisms that fuel this aberrant immune response and whether immune dysregulation is a cause or effect of these diseases. Another limitation is that our controls were relatively younger than the other conditions studied but this did not affect our analysis since we focused on the comparisons between COVID-19, post-COVID19-ILD and IPF. Finally, we did not have data regarding COVID-19 variants in these patients.

**Figure 5.**
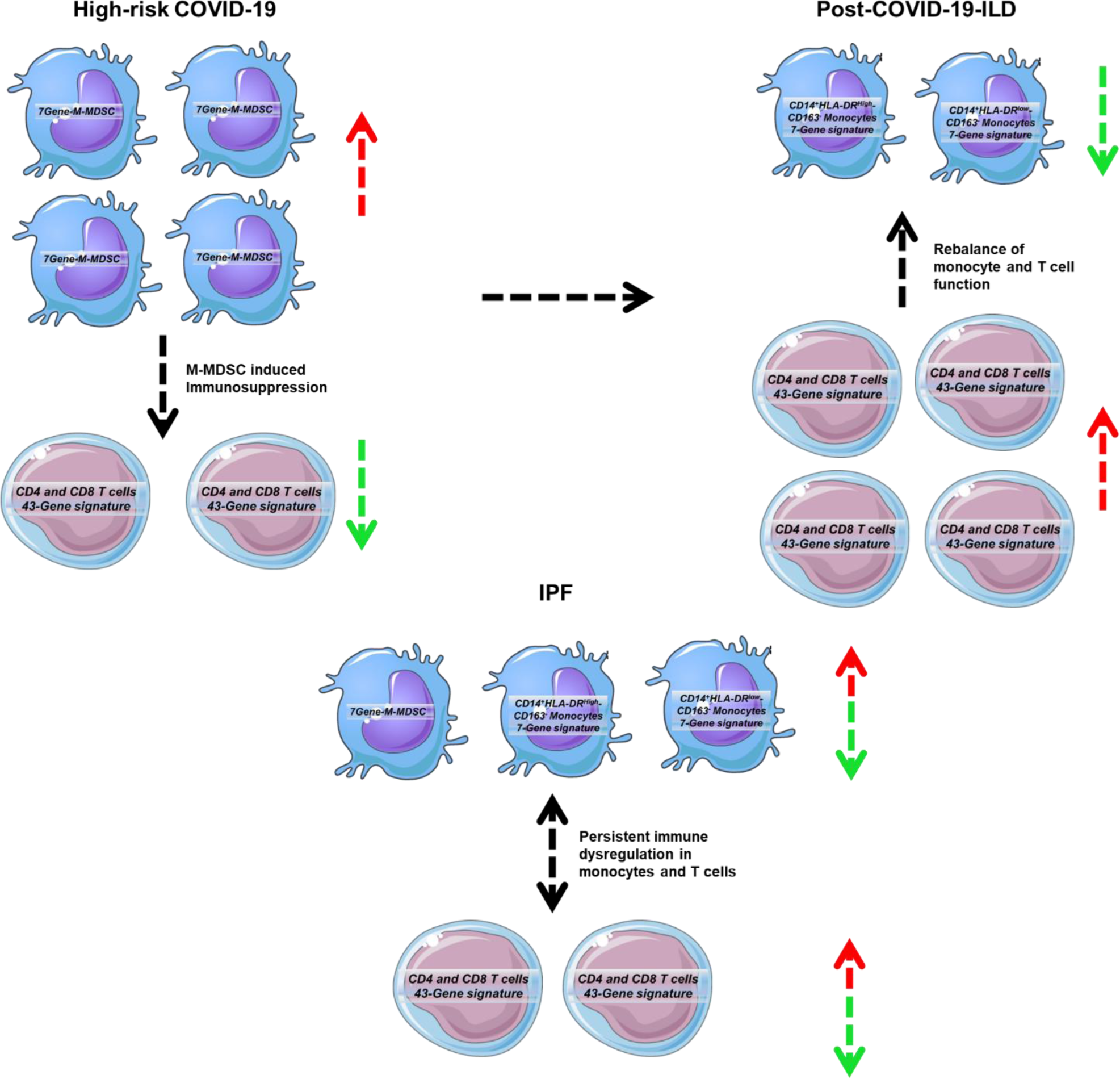
Potential model of changes in circulating immune cells expressing genes of the 50-gene signature in COVID-19, post-COVID-19-ILD and IPF. Our findings suggest that COVID-19 patients with a 50-gene high-risk profile have an imbalance between high 7Gene-M-MDSCs and low CD4 and CD8 T cell subsets that may be caused by the immunosuppressive effects that 7Gene-M-MDSCs exert in T cells. In post-COVID-19 ILD, the 7Gene-M-MDSCs transition to CD14^+^HLA-DR^high^CD163^−^ and CD14+HLA-DR^low^CD163^−^ leading to a recovery in the expression of genes of the 43-gene signature in CD4 and CD8 T cell subsets. In IPF patients, repeated cycles of alveolar epithelial cell injury sustain the presence of the subtypes of monocytes identified in our study, including 7Gene-M-MDSCs which leads to persistently low expression of genes of the 43-gene signature in T cell subsets.

Collectively, a 50-gene low-risk genomic profile in the peripheral blood was consistently predictive of COVID-19 survival. Comparison of this genomic profile through scRNA-seq in patients with COVID-19, post-COVID-19-ILD and IPF suggests an initial aberrant immune response in COVID-19 that resolves over time. This aberrant immune response is characterized by increased expression of 7Gene-M-MDSCs and decreased expression of CD4 T and CD8 T cell subsets. Survivors with post-COVID-19-ILD present with increased expression of T cell-related genes of the 43-gene signature compared to patients with COVID19 and IPF. This highlights that increased 7Gene-M-MDSCs and decreased T-cell subpopulations may have detrimental effects both in COVID-19 and IPF. Future studies looking at whether increased 7Gene-M-MDSCs and decreased T-cell subpopulation expressing genes of the 43-gene signature can be observed in patients with progressive forms of post-COVID-19-ILD and other forms of ILD are highly anticipated.

## Supplementary figures

**Supplementary figure 1:**
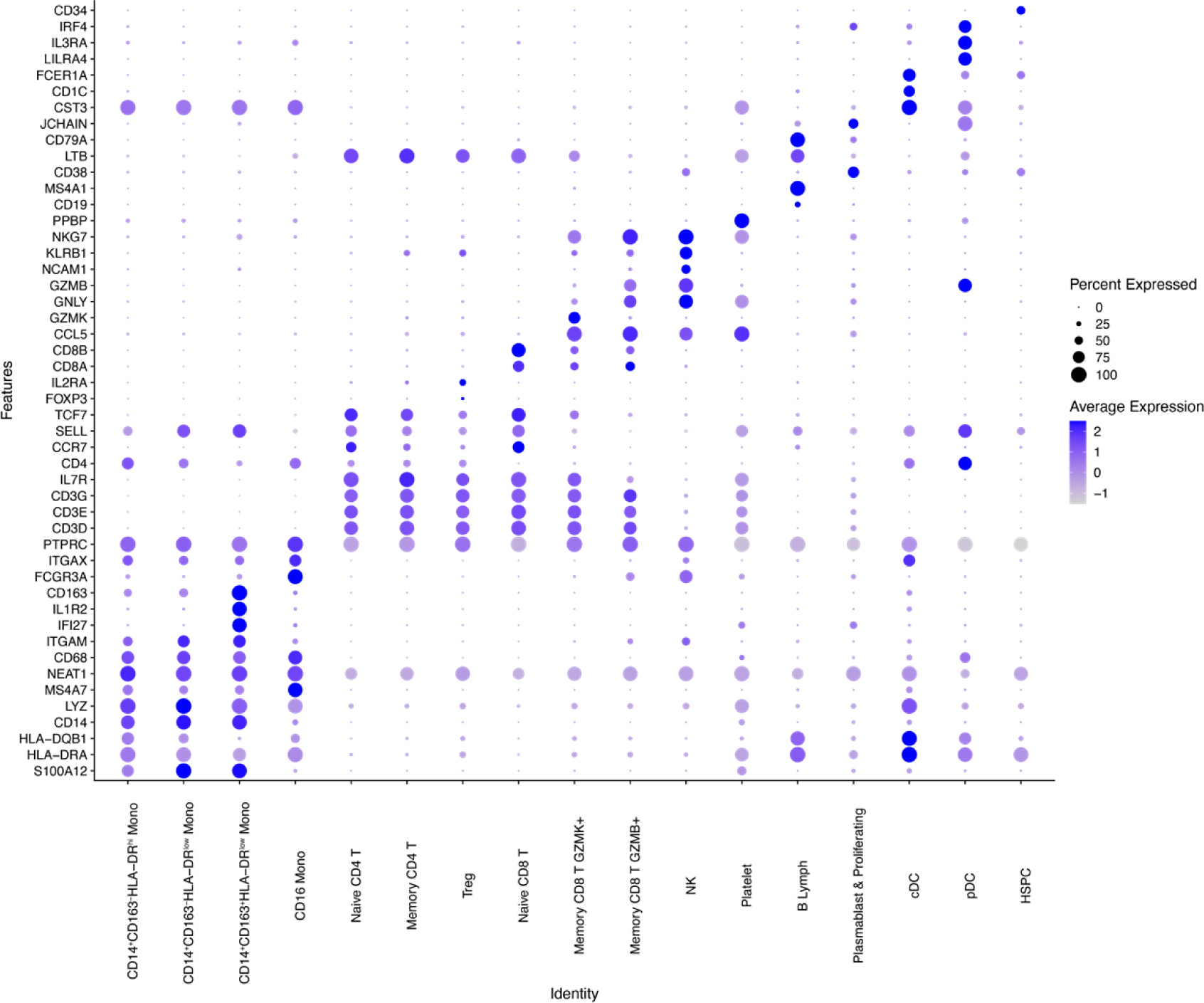
All immune cell clusters were identified based on the expression levels of different markers by reference to COVID-19 and IPF cell atlas. The Y axis represents markers and x axis represents the identified cells. Dot size is proportional to the percentage of cells expressing the gene in each subcluster. Color intensity is proportional to the average scaled log2-normalized expression within the cell subcluster.

**Supplementary figure 2:**
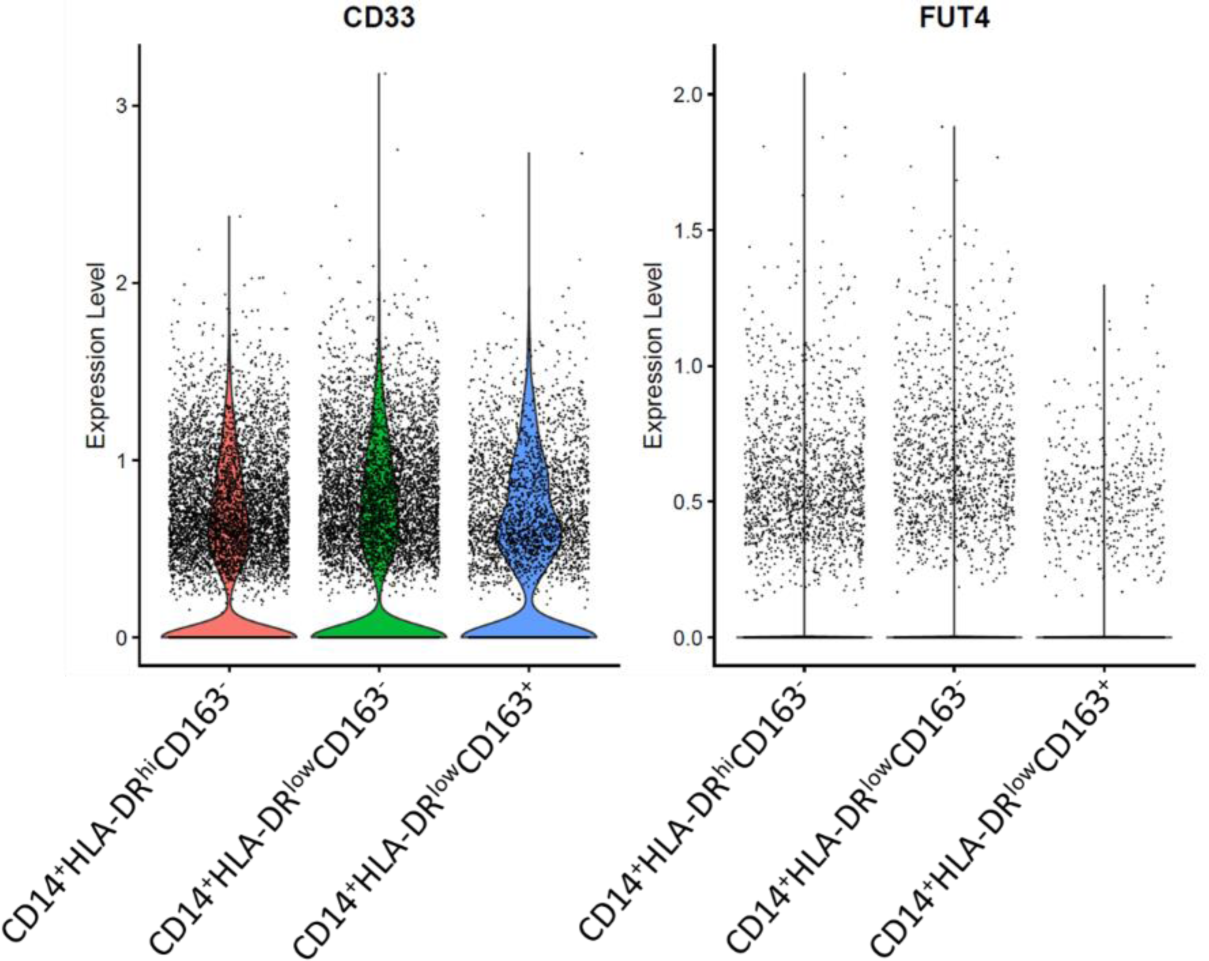
monocytes HLA-DR^low^ expressing CD33^+^ and CD15^−^ (FUT4) qualifying as Monocytic-Myeloid Derived Suppressive Cells (M-MDSCs), one of them (HLA-DR^low^CD163^+^), is expressed exclusively in COVID-19 patients.

